# Oligodendrocyte-lineage cell exocytosis and L-type prostaglandin D synthase promote oligodendrocyte development and myelination

**DOI:** 10.1101/2022.02.14.480339

**Authors:** Lin Pan, Amelia Trimarco, Alice J. Zhang, Ko Fujimori, Yoshihiro Urade, Lu O. Sun, Carla Taveggia, Ye Zhang

## Abstract

In the developing central nervous system, oligodendrocyte precursor cells (OPCs) differentiate into oligodendrocytes, which form myelin around axons. Oligodendrocytes and myelin are essential for the function of the central nervous system, as evidenced by the severe neurological symptoms that arise in demyelinating diseases such as multiple sclerosis and leukodystrophy. Although many cell-intrinsic mechanisms that regulate oligodendrocyte development and myelination have been reported, it remains unclear whether interactions among oligodendrocyte-lineage cells (OPCs and oligodendrocytes) affect oligodendrocyte development and myelination. Here, we show that blocking vesicle-associated membrane protein (VAMP) 1/2/3-dependent exocytosis from oligodendrocyte-lineage cells impairs oligodendrocyte development, myelination, and motor behavior in mice. Adding oligodendrocyte-lineage cell-secreted molecules to secretion-deficient OPC cultures partially restores the morphological maturation of oligodendrocytes. Moreover, we identified L-type prostaglandin D synthase as an oligodendrocyte-lineage cell-secreted protein that promotes oligodendrocyte development and myelination *in vivo*. These findings reveal a novel autocrine/paracrine loop model for the regulation of oligodendrocyte and myelin development.

## Introduction

In the developing central nervous system (CNS), oligodendrocyte precursor cells (OPCs) differentiate into oligodendrocytes (Bergles and Richardson, 2016; Hill et al., 2014; Kang et al., 2010), which form myelin sheaths around axons. Myelin is essential for the propagation of action potentials and for the metabolism and health of axons (Fünfschilling et al., 2012; Larson et al., 2018; Mukherjee et al., 2020; Saab et al., 2016; Schirmer et al., 2018; Simons and Nave, 2016). When oligodendrocytes and myelin are damaged in demyelinating diseases such as multiple sclerosis (MS) and leukodystrophy, sensory, motor, and cognitive deficits can ensue (Gruchot et al., 2019; Lubetzki et al., 2020; Stadelmann et al., 2019). In a broader range of neurological disorders involving neuronal loss, such as brain/spinal cord injury and stroke, the growth and myelination of new axons are necessary for neural repair (Wang et al., 2020). Thus, understanding oligodendrocyte development and myelination is critical for developing treatments for a broad range of neurological disorders.

Over the past several decades, researchers have made great progress in elucidating the cell-intrinsic regulation of oligodendrocyte development and myelination (e.g., transcription factors, epigenetic mechanisms, and cell death pathways) (Aggarwal et al., 2013; Bergles and Richardson, 2016; Budde et al., 2010; Dugas et al., 2010; Elbaz and Popko, 2019; Elbaz et al., 2018; Emery and Lu, 2015; Emery et al., 2009; Fedder- Semmes and Appel, 2021; Foerster et al., 2020; Harrington et al., 2010; Herbert and Monk, 2017; Howng et al., 2010; Koenning et al., 2012; Mitew et al., 2018; Nawaz et al., 2015; Snaidero et al., 2017; Sun et al., 2018; Wang et al., 2017; Xu et al., 2020; Zhao et al., 2018; Zuchero et al., 2015), as well as the cell-extrinsic regulation by other cell types (e.g., neurons (Gibson et al., 2014; Hines et al., 2015; Mayoral et al., 2018; Osso et al., 2021; Redmond et al., 2016; Wake et al., 2011), microglia/macrophages (Butovsky et al., 2006; Sherafat et al., 2021), and lymphocytes (Dombrowski et al., 2017)). However, it remains unclear whether interactions among oligodendrocyte-lineage cells (OPCs and oligodendrocytes) affect oligodendrocyte development and myelination.

One of the most abundant proteins secreted by oligodendrocyte-lineage cells is lipocalin-type prostaglandin D synthase (L-PGDS) (Zhang et al., 2014, 2016). Oligodendrocytes and meningeal cells are major sources of L-PGDS in the CNS (Urade et al., 1993; Zhang et al., 2014, 2016). L-PGDS has two functions: as an enzyme and as a carrier (Urade and Hayaishi, 2000). As an enzyme, L-PGDS converts prostaglandin H2 to prostaglandin D2 (PGD2). PGD2 regulates sleep, pain, and allergic reactions (Eguchi et al., 1999; Satoh et al., 2006; Urade and Hayaishi, 2011). L-PGDS also binds and transports lipophilic molecules such as thyroid hormone, retinoic acid, and amyloid-β (Urade and Hayaishi, 2000) and promotes Schwann cell myelination in the peripheral nervous system (Trimarco et al., 2014). Yet, its function in the development of the CNS is unknown.

To determine whether cell-cell interactions within the oligodendrocyte lineage regulate oligodendrocyte development, we blocked VAMP1/2/3-dependent exocytosis from oligodendrocyte-lineage cells *in vivo* and found impairment in oligodendrocyte development, myelination, and motor behavior in mice. Similarly, exocytosis-deficient OPCs exhibited impaired development *in vitro.* Adding oligodendrocyte-lineage cell-secreted molecules promoted oligodendrocyte development. These results suggest that an autocrine/paracrine loop promotes oligodendrocyte development and myelination. We assessed L-PGDS as a candidate autocrine/paracrine signal and further discovered that oligodendrocyte development and myelination were impaired in L-PGDS-knockout mice. Moreover, PGD2 partially restored the morphological maturation of exocytosis-deficient OPCs. Thus, L-PGDS is an oligodendrocyte-lineage cell-secreted protein that promotes oligodendrocyte development. These results reveal a new autocrine/paracrine loop model for the regulation of oligodendrocyte development in which VAMP1/2/3-dependent exocytosis from oligodendrocyte-linage cells and secreted L-PGDS promote oligodendrocyte development and myelination.

## Results

### Expression of botulinum toxin B in oligodendrocyte-lineage cells *in vivo*

If oligodendrocyte-lineage cells use autocrine/paracrine mechanisms to promote development and myelination, one would predict that (1) blocking secretion from oligodendrocyte-lineage cells would impair oligodendrocyte development and myelination and, in turn, that (2) adding oligodendrocyte-lineage cell-secreted molecules might promote oligodendrocyte development. Membrane fusion relies on soluble N-ethylmaleimide-sensitive fusion protein attachment protein receptors (SNARE) family proteins located on vesicles (v-SNAREs) and target membranes (t-SNAREs). Binding of v-SNAREs and t-SNARES form intertwined α-helical bundles that generate force for membrane fusion (Pobbati et al., 2006). VAMP1/2/3 are v-SNAREs that drive the fusion of vesicles with the plasma membrane to mediate exocytosis (Chen and Scheller, 2001). We found that oligodendrocyte-lineage cells express high levels of VAMP2 and VAMP3 and low levels of VAMP1 *in vivo* (Fig. 1A-C) (Zhang et al., 2014, 2016), consistent with previous reports *in vitro* (Feldmann et al., 2009, 2011; Madison et al., 1999). Botulinum toxin B specifically cleaves VAMP1/2/3 (Yamamoto et al., 2012), but not VAMP4, 5, 7, or 8 (Yamamoto et al., 2012), and inhibits the release of vesicles containing proteins (Somm et al., 2012) as well as small molecules such as neural transmitters (Poulain et al., 1988). Of note, botulinum toxin B does not cleave VAMP proteins that are involved in the vesicular transport between the trans-Golgi network, endosomes, and lysosomes (Antonin et al., 2000; Hoai et al., 2007; Pols et al., 2013). Similarly, botulinum toxins do not affect ion channel- or membrane transporter-mediated release of small molecules.

**Figure 1.**
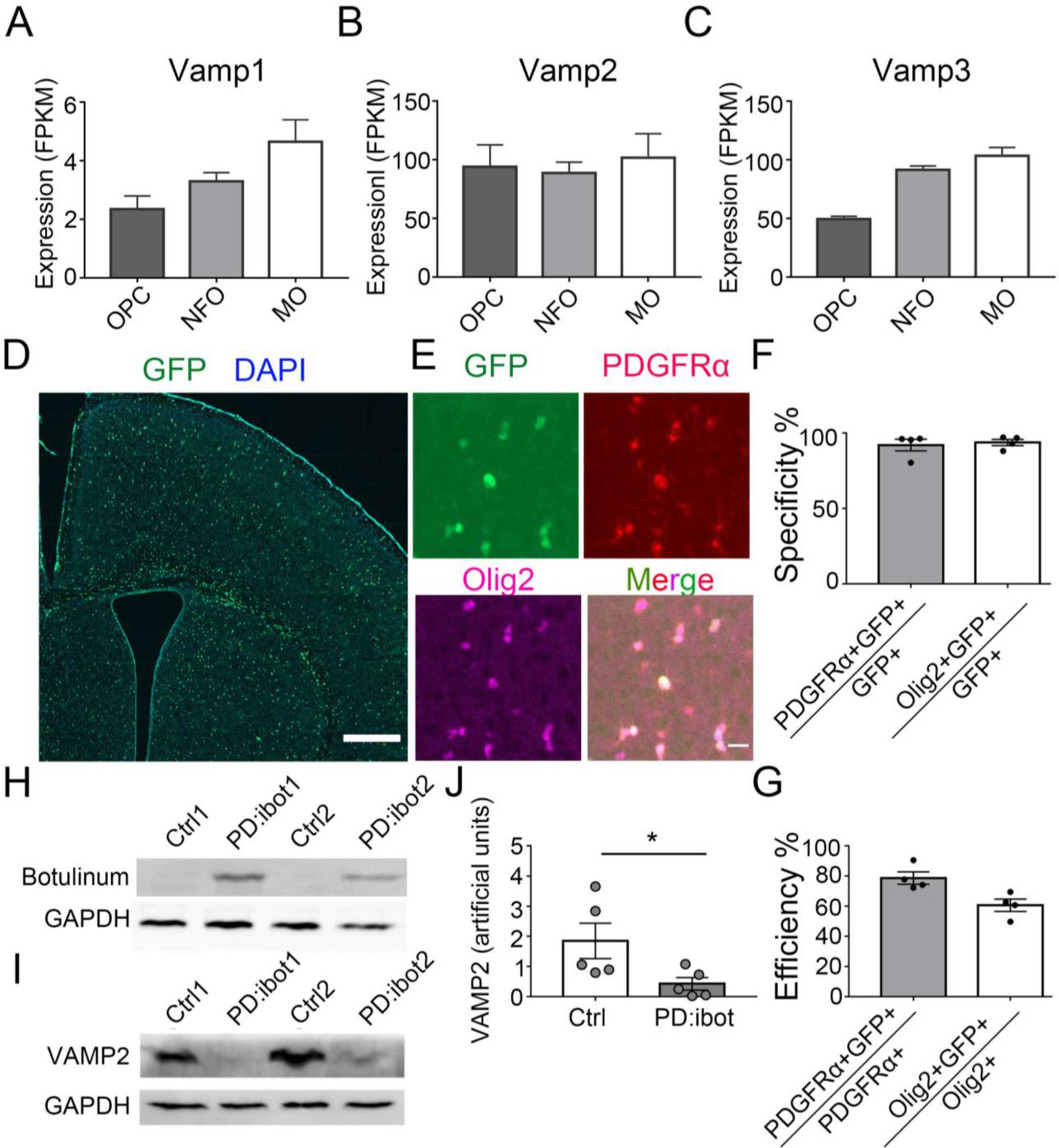
VAMP1/2/3 and ibot expression in oligodendrocyte-lineage cells. (A-C) Expression of VAMP1/2/3 by oligodendrocyte-lineage cells determined by RNA-seq (Zhang et al., 2014). NFO, newly formed oligodendrocytes. MO, myelinating oligodendrocytes. (D) Expression of ibot-GFP in PDGFRα-CreER; ibot (PD:ibot) mice. Scale bar: 500 μm. (E) Colocalization of ibot-GFP with PDGFRα and Olig2 in PD:ibot mice. Scale bar: 20 μm. (F) Specificity of ibot-GFP expression in oligodendrocyte-lineage cells. N=4 mice per group. 92.1±3.9% of GFP^+^ cells were PDGFRα^+^; 93.8±2.1% of GFP^+^ cells were Olig2^+^. (G) Efficiency of ibot-GFP expression in oligodendrocyte-lineage cells. N=4 mice per group. 78.7±4.1% of PDGFRα^+^ cells were GFP^+^; 60.6±4.1% of Olig2^+^ cells were GFP^+^. P8 mice were used in (D-G). (H) Presence of botulinum toxin B-light chain in oligodendrocyte cultures from 4- hydroxytamoxifen-injected PD:ibot mice detected by Western blots. (I) Reduced levels of full-length VAMP2 in oligodendrocyte cultures from 4- hydroxytamoxifen-injected PD:ibot mice determined by Western blots. (J) Quantification of VAMP2 immunoblot signal intensity. N=5 mice per group. Paired two-tailed T-test. *, p<0.05. **, p<0.01. ***, p<0.001. NS, not significant.

To block VAMP1/2/3-dependent exocytosis from oligodendrocyte-lineage cells, we crossed PDGFRα-CreER transgenic mice, which express Cre recombinase in OPCs (PDGFRα^+^Olig2^+^) (Kang et al., 2010), with loxP-stop-loxP-botulinum toxin B light chain- IRES-green fluorescent protein (GFP) (inducible botulinum toxin B, or ibot) transgenic mice (Slezak et al., 2012), allowing expression of botulinum toxin B-light chain in OPCs and their progeny. The light chain contains the catalytically active domain of the toxin but lacks the heavy chain, which allows cell entry (Montal, 2010), thus confining toxin expression to the targeted cell type. Therefore, the ibot transgenic mice allow for the inhibition of VAMP1/2/3-dependent exocytosis in a cell-type-specific and temporally controlled manner (Slezak et al., 2012).

In our study, we used double-transgenic mice hemizygous for both Cre and ibot and referred to them as the PD:ibot mice thereafter. To validate our model and test its recombination efficiency, we injected 0.1 mg of 4-hydroxytamoxifen in each PD:ibot mouse daily for 2 days between postnatal day 2-4 (P2-4) and examined GFP expression at P8 and P30. We assessed whether GFP expression is restricted to oligodendrocyte-lineage cells (specificity) and what proportion of oligodendrocyte-lineage cells express GFP (efficiency/coverage). At P8, when the vast majority of oligodendrocyte-lineage cells are undifferentiated OPCs, we detected specific expression of GFP in oligodendrocyte-lineage cells (Fig. 1D-F). GFP was efficiently expressed by oligodendrocyte-lineage cells (Fig. 1G). At P30, when substantial numbers of OPCs have differentiated (PDGFRα^-^ Olig2^+^), we observed a similarly high specificity of GFP expression in oligodendrocyte-lineage cells (Figure 1-figure supplement 1). These observations are consistent with previous reports on the specificity and efficiency of the PDGFRα-CreER transgenic line (Kang et al., 2010). As controls, we used mice with only the Cre transgene or only the ibot transgene subjected to the same tamoxifen injection scheme. In both control conditions, we detected very little GFP expression.

To directly assess the expression of botulinum toxin B-light chain and the cleavage of the VAMP proteins in oligodendrocyte-lineage cells from PD:ibot mice, we purified OPCs from PD:ibot and control mice by immunopanning and allowed them to differentiate into oligodendrocytes in culture. We performed Western blot analysis of the cultures and detected botulinum toxin B-light chain in PD:ibot but not in control cells (Fig. 1H).

Furthermore, levels of full-length VAMP2 proteins were lower in PD:ibot cells compared with control cells (Fig. 1I, J). Based on these observations, we conclude that the botulinum toxin-GFP transgene is specifically and efficiently expressed by oligodendrocyte-lineage cells in PD:ibot mice.

### Blocking VAMP1/2/3-dependent exocytosis from oligodendrocyte-lineage cells impairs oligodendrocyte development, myelination, and motor behavior

In PD:ibot mice, we found that the numbers of differentiated oligodendrocytes (CC1^+^) were reduced in the cerebral cortex (Fig. 2A, B), whereas the number of OPCs (PDGFRα^+^) did not change (Fig. 2C, E, F, H). Olig2 labels both OPCs and differentiated oligodendrocytes, and the densities of Olig2^+^ cells in PD:ibot mice are also reduced, likely due to the reduction in differentiated oligodendrocytes (Fig. 2C, D, F, G). At P8, the vast majority of PDGFRα-CreER-expressing cells are OPCs (Paukert et al., 2014). Therefore, it is more likely that blocking exocytosis from OPCs rather than oligodendrocytes affects oligodendrocyte development during the early postnatal period.

**Figure 2.**
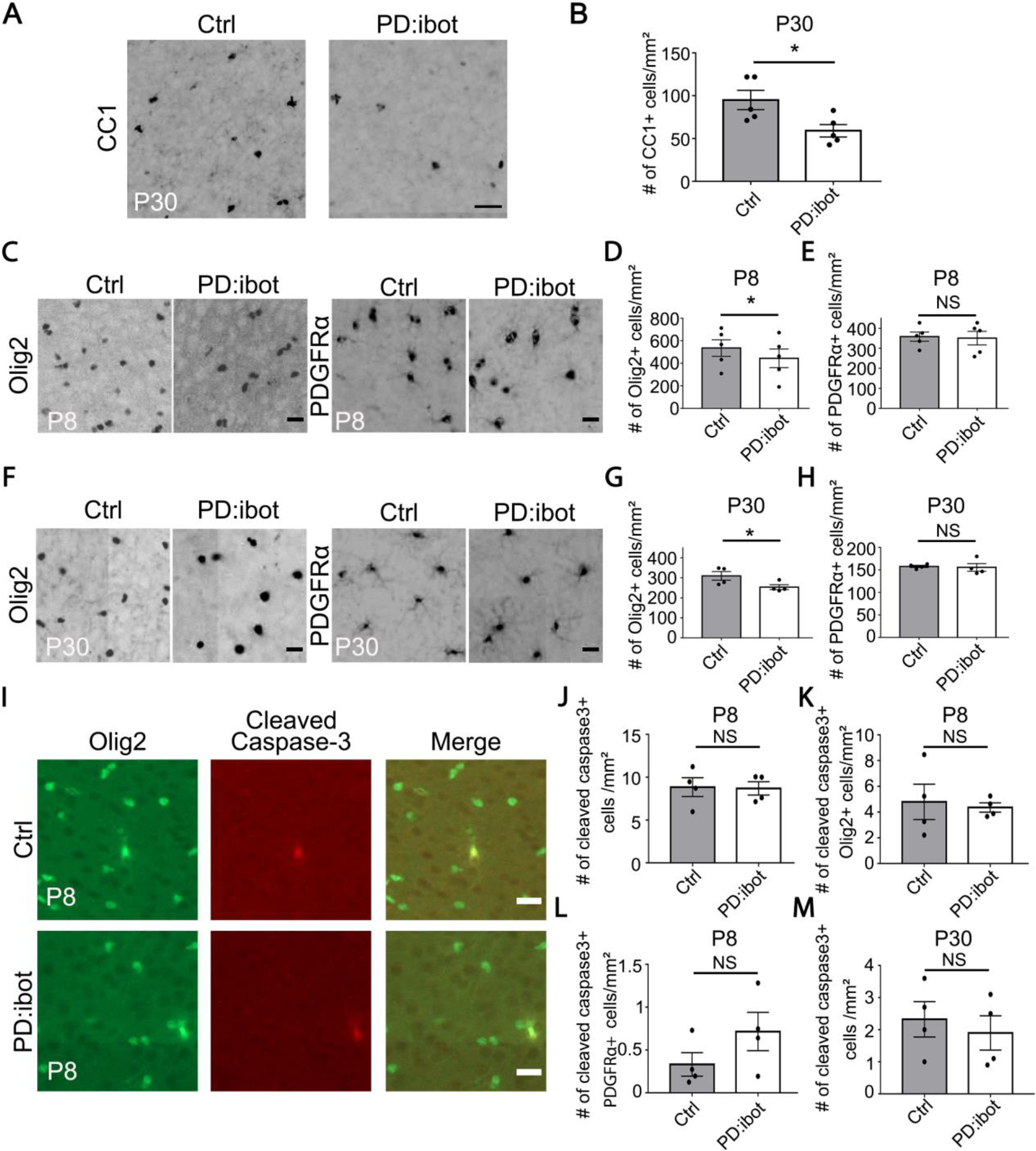
Reduction of CC1^+^ and Olig2^+^ oligodendrocytes in PD:ibot mic. (A) Differentiated oligodendrocytes labeled by CC1 in the cerebral cortex of PD:ibot and control mice at P30. Scale bar: 50 μm. (B) Quantification of the density of CC1^+^ differentiated oligodendrocytes in the cerebral cortex of PD:ibot and control mice at P30. N=5 mice per group. Paired two-tailed T-test. CC1^+^ cells/mm^2^: 95.04±11.22 in control and 59.04±7.28 in PD:ibot, p=0.049. (C, F) Olig2^+^ oligodendrocyte-lineage cells and PDGFRα^+^ OPCs in the cerebral cortex of PDibot and control mice at P8 (C) and P30 (F). Scale bars: 20 μm. (D, E, G, H) Quantification of the density of Olig2^+^ and PDGFRα^+^ cells in the cerebral cortex of PD:ibot and control mice at P8 and P30. N=5 mice per group at P8. N=4 mice per group at P30. Paired two-tailed T-test. Olig2^+^ cells/mm^2^: 535.6±73.6 in control and 444.4±82.9 in PD:ibot, p=0.040 at P8; 309.9±21.7 in control and 253.8±12.2 in PD:ibot, p=0.048 at P30. PDGFRα^+^ cells/mm^2^: 358.4±22.8 in control and 350.9±34.0 in PD:ibot, p=0.74 at P8; 157.8±2.6 in control and 155.6±8.5 in PD:ibot, p=0.85 at P30. (I) Examples of apoptotic oligodendrocyte-lineage cells labeled by cleaved caspase-3 in the cerebral cortex at P8. Scale bar: 20 μm. (J-M) Quantification of the density of caspase3^+^ cells in the cerebral cortex of PD:ibot and control mice. N=4 mice per group. Paired two-tailed T-test. P8: Caspase3^+^ cells/mm^2^: 8.9±1.1 in control and 8.7±0.8 in PD:ibot; p=0.93; Caspase3^+^PDGFRα^+^ cells/mm^2^: 0.33±0.14 in control and 0.72±0.22 in PD:ibot; p=0.33; Caspase3^+^Olig2^+^ cells/mm^2^: 4.8±1.4 in control and 4.4±0.4 in PD:ibot; p=0.74. P30: Caspase3^+^ cells/mm^2^: 2.3±1.1 in control and 1.9±1.1 in PD:ibot; p=0.53.

We next examined myelin development in PD:ibot mice and found that immunofluorescence of myelin basic protein (MBP), one of the main components of CNS myelin, is reduced in PD:ibot mice (Fig. 3A-L). Moreover, many MBP^+^ ibot-GFP- expressing cells exhibit round cell morphology whereas MBP^+^GFP^-^ control cells form elongated myelin internodes along axon tracks (Fig. 3M). Transmission electron microscopy allows for the assessment of myelin structure at a high resolution. Thus, to further examine myelination in PD:ibot mice, we performed transmission electron microscopy imaging and found a reduction in the percentage of myelinated axons (Fig. 3N, O) and reduced myelin thickness in PD:ibot mice (g-ratio: axon diameter divided by the diameter of axon + myelin; Fig. 3P, Q).

**Figure 3.**
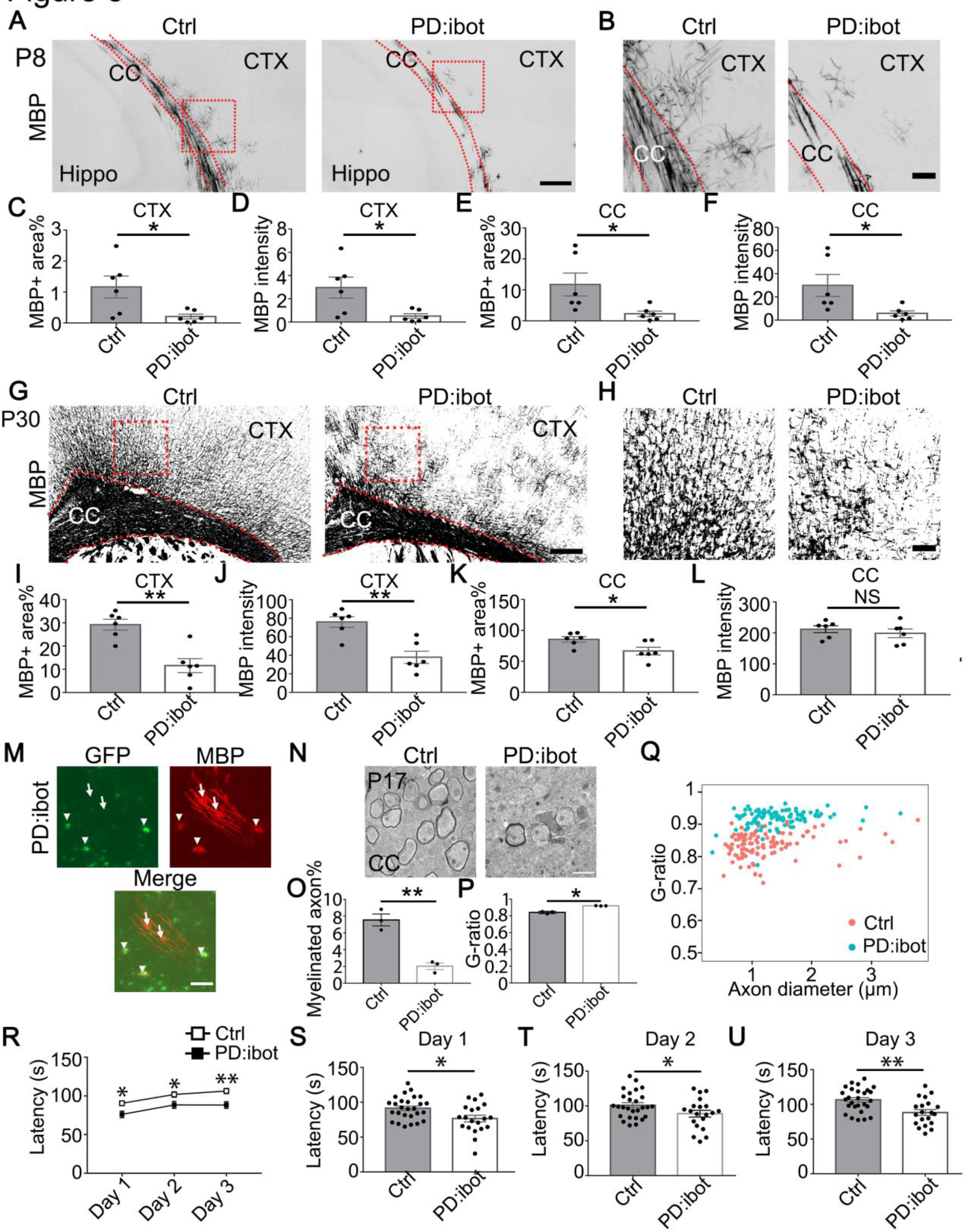
Defective myelination and motor behavior in PD:ibot mice. (A, B) MBP immunofluorescence at P8 in PD:ibot and control brains. Dashed lines delineate corpus callosum. CTX, cerebral cortex. CC, corpus callosum. Hippo, hippocampus. Boxed areas in (A) are enlarged and shown in (B). Scale bars: 200 μm in (A), 50 μm in (B). (C-F) Quantification of MBP^+^ area and mean MBP fluorescence intensity in the cerebral cortex (C, D) and the corpus callosum (E, F) at P8. N=6 mice per group. Paired two-tailed T-test. (G, H) MBP immunofluorescence at P30 in PD:ibot and control brains. Dashed lines delineate corpus callosum. Boxed areas in (G) are enlarged and shown in (H). Scale bars: 200 μm in (G), 50 μm in (H). (I-L) Quantification of MBP^+^ area and mean MBP fluorescence intensity in the cerebral cortex (I, J) and corpus callosum (K, L) at P30. N=6 mice per group. Paired two-tailed T-test. (M) The morphology of ibot-GFP^+^ cells and GFP^-^ control cells labeled by MBP immunofluorescence. A region in the cerebral cortex from a P8 PD:ibot mouse is shown. The arrowheads point to ibot-GFP^+^ cells and the arrows point to GFP^-^ control cells. Scale bar: 50 μm. (N) Transmission electron microscopy images of the corpus callosum at P17. Scale bar: 1 μm. (O, P) Quantification of the percentage of myelinated axons and G-ratio (axon diameter divided by the diameter of myelin + axon) from the transmission electron microscopy images of the corpus callosum at P17. N=3 mice per group. Paired two-tailed T-test. Myelinated axons %: 7.6±0.7 in control and 2.0±0.4 in PD:ibot, p=0.0048. G-ratio: 0.84±0.009 in control and 0.92±0.0009 in PD:ibot; p=0.012. (Q) G-ratio as a function of axon diameter in the corpus callosum at P17. N=3 mice per group. (R) Latency to fall from an accelerating rotarod (seconds). Each mouse was tested three times per day for three consecutive days. The average latency to fall of the three trials of each mouse was recorded for each day. No significant sex differences were detected. Unpaired two-tailed T-test was performed with Benjamini, Krieger, and Yekutieli’s false discovery rate (FDR) method to correct for multiple comparisons. *, FDR <0.05. **, FDR<0.01. (S-U) Latency to fall on each testing day. Day 1: PD:ibot: 76.7±4.7 seconds, control: 91.4±3.2 seconds, p=0.015; day 2: PD:ibot: 88.9±4.9 seconds, control: 101.1±3.7 seconds, p=0.033; day 3: PD:ibot: 88.4±4.3 seconds, control: 106.4±3.3 seconds, p=0.0051. N=27 mice for control and 20 mice for PD:ibot.

To determine whether the reduction of oligodendrocytes in PD:ibot mice is caused by cell death, we performed immunostaining with an antibody against activated caspase- 3, which labels apoptotic cells. We observed no difference in the total apoptotic cells (caspase-3^+^), apoptotic OPCs (caspase-3^+^PDGFRα^+^), or apoptotic oligodendrocyte- lineage cells (caspase-3^+^Olig2^+^) between PD:ibot and control mice in the cerebral cortex (Fig. 2I-M).

To investigate whether the expression of botulinum toxin B-light chain affects oligodendrocyte development and myelination in non-cell-type-specific manners, we blocked exocytosis from astrocytes or endothelial cells by crossing ibot transgenic mouse with mGFAP-Cre (line 77.6) and Tie2-Cre strains, respectively. We found astrocyte- and endothelial cell-specific expression of ibot-GFP in these mice but did not detect any obvious changes in oligodendrocyte density or myelin proteins (data not shown). These observations suggest that botulinum toxin B-light chain peptides have specific effects on the targeted cell types.

To examine the functional consequences of blocking VAMP1/2/3-dependent exocytosis from oligodendrocyte-lineage cells, we assessed the motor behavior of PD:ibot and littermate control mice using the rotarod test. We placed mice on a gradually accelerating rotarod and recorded the time each mouse stayed on the rotarod. We found that PD:ibot mice stayed on the rotarod for significantly shorter amounts of time than littermate control mice on all three days of testing (Fig. 3R-U). Therefore, blocking VAMP1/2/3-dependent exocytosis from oligodendrocyte-lineage cells led to deficits in neural circuit function.

### Blocking VAMP1/2/3-dependent exocytosis from oligodendrocyte-lineage cells impairs oligodendrocyte development *in vitro*

VAMP1/2/3-dependent exocytosis from oligodendrocyte-lineage cells may directly affect oligodendrocyte development or change the attributes of other cell types, and, in turn, indirectly affect oligodendrocytes. For example, OPC-secreted molecules may affect axonal growth, and subsequently axonal signals may affect oligodendrocytes indirectly. Therefore, we next employed purified OPC and oligodendrocyte cultures to determine whether secreted molecules have direct autocrine/paracrine roles in oligodendrocyte-lineage cells in the absence of other cell types.

We performed immunopanning to purify OPCs from P7 PD:ibot and control mice injected with 4-hydroxytamoxifen as described above. We cultured the OPCs for two days in the proliferation medium and then switched to the differentiation medium and cultured them for another seven days. To assess oligodendrocyte differentiation and maturation, we assessed the levels of MBP protein, a marker for differentiated oligodendrocytes, and detected lower MBP levels in PD:ibot cells compared with controls (Fig. 4B, C).

**Figure 4.**
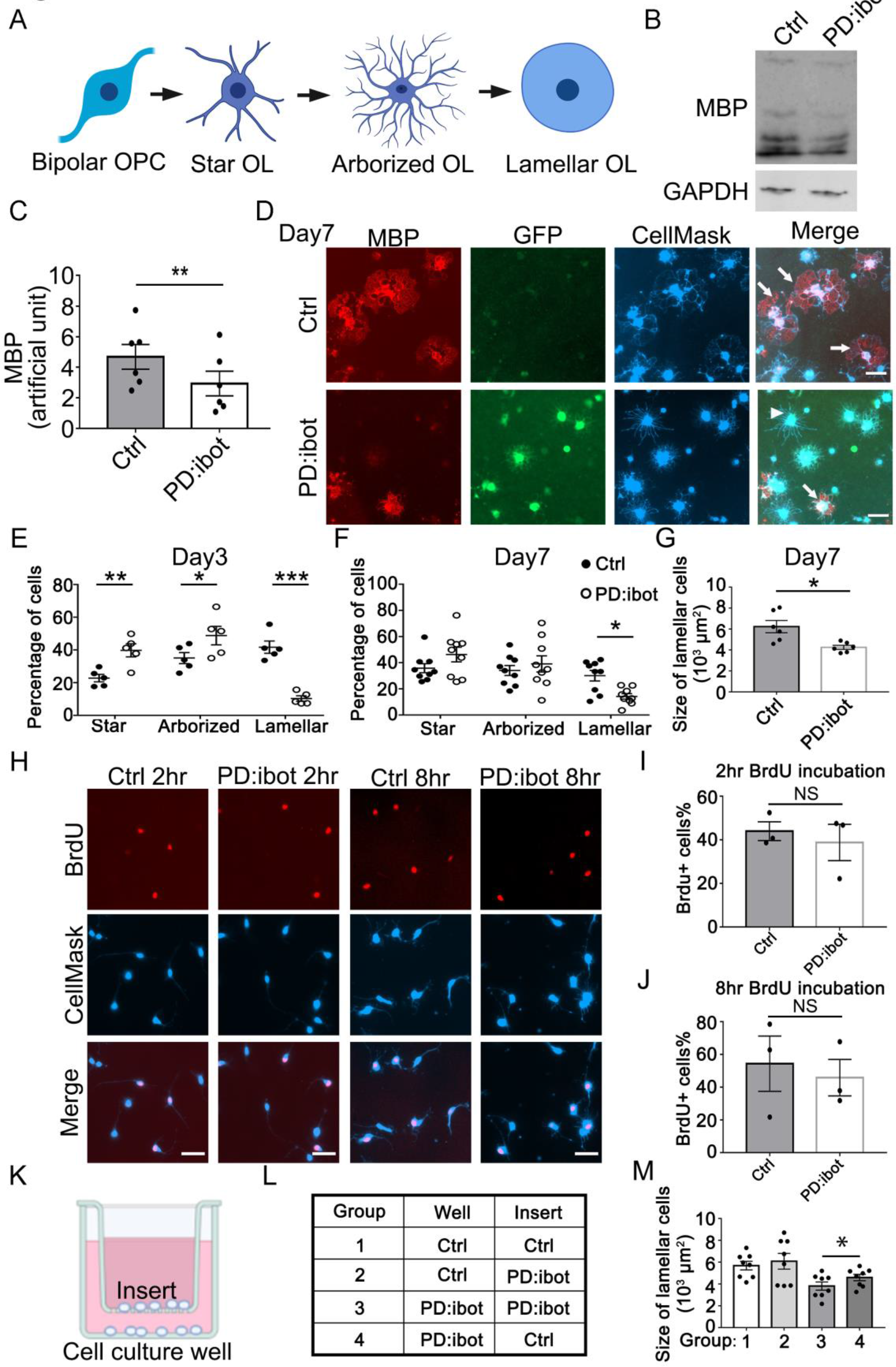
*In vitro* development defect of oligodendrocytes purified from PD:ibot mice. (A) A diagram of the morphological changes during oligodendrocyte differentiation *in vitro*. OPCs exhibit a bipolar morphology. Differentiating oligodendrocytes first grow multiple branches (star-shaped and arborized) and then develop myelin-like membrane extension and exhibit a lamellar morphology. (B) Western blot for MBP in oligodendrocyte cultures after 7 days of differentiation. (C) Quantification of MBP proteins from Western blot. N=6 mice per group. Paired two-tailed T-test. (D) Oligodendrocyte cultures after 7 days of differentiation. Red, MBP. Green, ibot:GFP. Blue, CellMask, which labels all cells. Arrows point to examples of lamellar cells and an arrowhead points to an example of a star-shaped oligodendrocyte. Scale bars: 50 μm. (E-F) Quantification of the percentage of cells at the star-shaped, arborized, and lamellar stages after 3 (E) and 7 (F) days of differentiation. Filled circles, control. Open circles, PD:ibot. N=5 mice per group on day 3 and N=9 mice per group on day 7. Unpaired two-tailed T-test with the Holm-Sidak multiple comparison adjustment. (G) Quantification of the size of lamellar cells in oligodendrocyte cultures obtained from PD:ibot and littermate control mice after 7 days of differentiation. N=6 cultures from 4 mice per group. Paired two-tailed T-test. 6,237±587.5 μm^2^ in control and 4,253±193.7 μm^2^ in PD:ibot; p=0.016. (H) Proliferation of OPCs from PD:ibot and littermate control mice *in vitro*. OPC cultures were fixed and stained 2 and 8 hours after the addition of BrdU. Hr, hour. Scale bars: 50 μm. (I, J) Quantification of the percentage of BrdU^+^ cells in OPC cultures obtained from PD:ibot and littermate control mice at 2 (I) and 8 (J) hours after addition of BrdU. N=3 mice per group. Paired two-tailed T-test. 2 hour BrdU incubation, BrdU^+^ cells%: 44.0±4.3 in control and 38.8±8.3 in PD:ibot; p=0.51; 8 hour BrdU incubation, BrdU^+^ cells%: 54.4±16.9 in control and 45.9±11.1 in PD:ibot; p=0.49. (K) Diagram of cocultures of cells separated by a porous insert with 1 μm pore size. (L) The genotype of cells on the inserts and wells in each group. The differentiation of the cells on the bottom of the wells was examined. N=8 cultures from 5 mice per condition. (M) Quantification of the size of lamellar cells. Group 3 *vs.* group 4: p=0.016. Group 1 *vs.* group 3: p=0.0002. Group 2 *vs.* group 3: p=0.0051. Group 1 *vs.* group 4: p=0.0068. All other pairs of conditions are not significantly different. One-way ANOVA with Tukey’s test for multiple comparisons. Size of lamellar cells: group 1: 5,691±391 μm^2^; group 2: 6,087±720.7 μm^2^; group 3: 3,810±376 μm^2^; group 4: 4,594±293.3 μm^2^.

Additionally, we assessed the morphological maturation of oligodendrocytes *in vitro* (Fig. 4C-G). OPCs are initially bipolar, and as they differentiate, they grow a few branches to become star-like. The cells next grow more branches to become arborized and then extend myelin-sheath-like flat membranous structures, acquiring a “lamellar” morphology (Fig. 4A) (Zuchero et al., 2015) (also referred to as a “fried egg” or “pancake” morphology). We used the CellMask dye that labels plasma membrane to analyze the morphological maturation of oligodendrocytes. At day 3 of differentiation, we found that a larger proportion of PD:ibot cells than control cells are at the early “star” stage whereas a smaller proportion of PD:ibot cells than control cells have proceeded to the late “lamellar” stage (Fig. 4E). At day 7 of differentiation, more PD:ibot cells have proceeded from the “star” to the “arborized” stage compared with day 3, but the percentage of cells that have proceeded to the late “lamellar” stage remains lower in PD:ibot than in control cultures (Fig. 4D, F). We next quantified the size of lamellar cells, which have large sheaths of myelin-like membrane. Interestingly, we found that lamellar cells from PD:ibot mice are significantly smaller than those from control mice (Fig. 4D, G). Together, these observations suggest that VAMP1/2/3-dependent exocytosis is required for the morphological maturation of oligodendrocytes and that oligodendrocyte-lineage cell-secreted molecules act directly on cells within the oligodendrocyte lineage to promote their development.

We next examined whether blocking VAMP1/2/3-dependent exocytosis affects OPC proliferation using bromodeoxyuridine (BrdU) pulse-chase experiments and did not observe any differences between cultured OPCs from PD:ibot and littermate control mice (Fig. 4H-J).

### Oligodendrocyte-lineage cell-secreted molecules partially restore oligodendrocyte morphological maturation in secretion-deficient cells

Having established the necessity of VAMP1/2/3-dependent exocytosis from oligodendrocyte-lineage cells for oligodendrocyte development and myelination, we next assessed whether adding oligodendrocyte-lineage cell-secreted molecules could restore differentiation in VAMP1/2/3-dependent exocytosis-deficient OPCs. We prepared co-cultures of OPCs separated by inserts with 1 μm-diameter pores to allow for the diffusion of secreted molecules (Fig. 4K). We plated PD:ibot and control cells on inserts and on the bottom of culture wells in four combinations: (1) control-inserts-control-wells; (2) PD:ibot- inserts-control-wells; (3) PD:ibot-inserts-PD:ibot-wells; and (4) control-inserts-PD:ibot- wells (Fig. 4L) and examined oligodendrocyte morphological differentiation on the bottom of culture wells by quantifying lamellar cells as described above. Comparing group 3 vs. group 4, we found that adding secreted molecules from control cells on inserts partially rescued the size of lamellar cells of PD:ibot cells on the bottom of culture wells (Fig. 4M), lending further support to the hypothesis that oligodendrocyte-lineage cell-secreted molecules promote oligodendrocyte development.

### PD:ibot mice exhibit changes in the transcriptomes of OPCs and oligodendrocytes

We next aimed to uncover the molecular changes in OPCs and oligodendrocytes in PD:ibot mice, and to identify candidate secreted molecules that regulate oligodendrocyte differentiation and myelination. We performed immunopanning to purify OPCs and oligodendrocytes from the brains of P17 PD:ibot and littermate control mice and performed RNA-sequencing (RNA-seq). We detected broad and robust gene expression changes in oligodendrocytes, and moderate changes in OPCs (Fig. 5A, B, Supplementary Table 1, 2), demonstrating that VAMP1/2/3-dependent exocytosis from oligodendrocyte-lineage cells is critical for establishing and/or maintaining the normal molecular attributes of oligodendrocytes and OPCs. Notably, the expression of signature genes of differentiated oligodendrocytes such as *Plp1, Mbp, Aspa, and Mobp* were significantly reduced in oligodendrocytes purified from PD:ibot mice compared with controls (Fig. 5E-H, Supplementary Table 1, 2). This result was not secondary to a reduction in oligodendrocyte density, as we loaded a similar amount of cDNA libraries from PD:ibot and control oligodendrocytes for sequencing, and processed all sequencing data with the same pipeline. Therefore, VAMP1/2/3-dependent exocytosis from oligodendrocyte-lineage cells is critical for the expression of mature oligodendrocyte genes. We next performed gene ontology (GO) analysis to reveal the molecular pathways and cellular processes altered in each type of glial cell in PD:ibot mice (Fig. 5I-L, Supplementary Table 3). Genes associated with the filopodium assembly, calcium ion transport, and plasma membrane raft assembly pathways were increased and genes associated with lipid biosynthetic process, axon ensheathment, and myelination pathways were reduced in oligodendrocytes in PD:ibot mice. Genes associated with the trans-synaptic signaling, chemical synaptic transmission, and GPCR signaling pathways were increased in OPCs in PD:ibot mice. To assess whether oligodendrocyte-lineage cell exocytosis affects other glial cell types, such as astrocytes and microglia, we also purified these cells by immunopanning and performed RNA-seq. We observed moderate changes in astrocytes and microglia (Fig. 5C, D). For example, genes associated with phagocytosis, such as CD68 and C1qc, were increased in microglia from PD:ibot mice (Supplementary Table 2), suggesting the importance of oligodendrocyte-secreted molecules in oligodendrocyte-microglial interactions.

**Figure 5.**
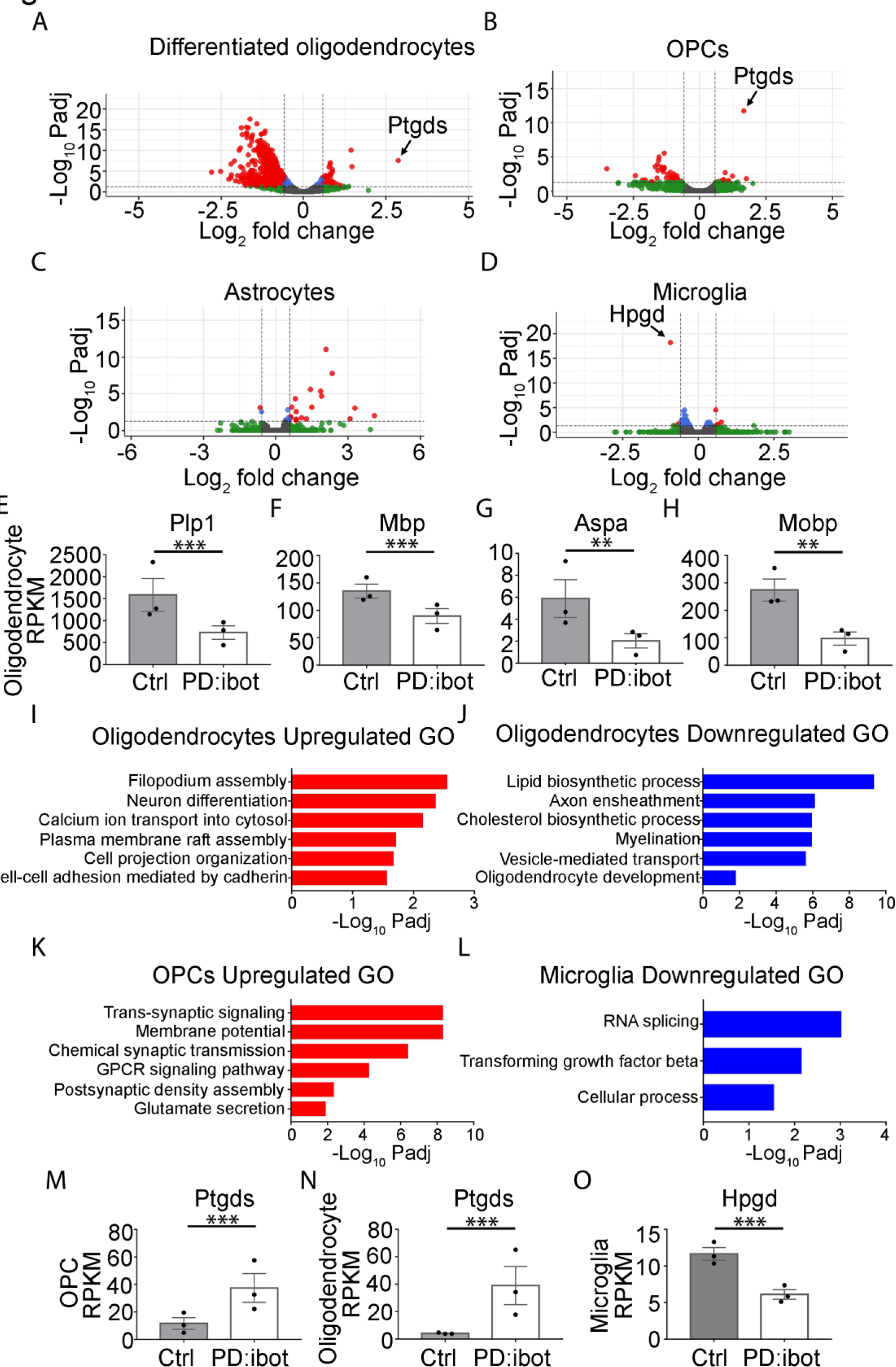
Transcriptome changes of purified glial cells from PD:ibot mice. (A-D) Differentiated oligodendrocytes, OPCs, astrocytes, and microglia were purified by immunopanning from whole brains of P17 PD:ibot and littermate control mice. Gene expression was determined by RNA-seq. Genes exhibiting significant changes (P-value adjusted for multiple comparisons, Padj <0.05, fold change >1.5) are shown in red. (E-H) Examples of mature oligodendrocyte marker gene expression by oligodendrocytes purified from PD:ibot and control mice at P17. (I-L) Examples of GO terms associated with genes upregulated in oligodendrocytes (I), downregulated in oligodendrocytes (J), upregulated in OPCs (K), and downregulated in microglia (L). There are no GO terms significantly associated with genes downregulated in OPCs or upregulated in microglia. (M, N) Expression of *Ptgds* by OPCs and oligodendrocytes purified from PD:ibot and control mice at P17. (O) Expression of *Hpgd* by microglia purified from PD:ibot and control mice at P17. (E-H, M-O) Expression is shown in RPKM. N=3 mice per group. Significance is determined by DESeq2.

We next sought to uncover the identity of the secreted molecules that promote oligodendrocyte and myelin development. If the signal is a protein, we posit that (1) oligodendrocyte-lineage cells must highly express the gene encoding the protein; (2) the expression pattern must be conserved in evolution in humans and mice; and (3) blocking its secretion should activate compensatory mechanisms to increase the expression of this gene and/or change the expression of related genes in the same pathway. We previously purified oligodendrocyte-lineage cells by immunopanning from human and mouse brains and performed RNA-seq (Zhang et al., 2014, 2016). We mined the data and generated a list of candidate genes meeting criteria 1 and 2. To identify genes meeting criterion 3, we analyzed the RNA-seq data from purified OPCs and oligodendrocytes from PD:ibot and littermate control mice. We found that the gene *Ptgds* (encoding the L-PGDS protein, which converts prostaglandin H2 to PGD2 (Urade and Hayaishi, 2000)) is highly upregulated specifically in OPCs and oligodendrocytes of PD:ibot mice (Fig. 5A, B, M, N), but not in microglia or astrocytes. *Ptgds* is one of the most abundant genes encoding a secreted protein expressed by oligodendrocyte-lineage cells in both humans and mice (Zhang et al., 2014, 2016). Its expression increases during development (Kang et al., 2011), as oligodendrocyte development and myelination occur, and is reduced in OPCs in multiple sclerosis patients (Jäkel et al., 2019). L-PGDS is also important for Schwann cell myelination in the peripheral nervous system (Trimarco et al., 2014). Yet, its function in the development of the CNS is unknown.

Of note, in microglia from PD:ibot mice, we found significant downregulation of the gene *Hpgd*, which encodes a PGD2 degradation enzyme (Conner et al., 2001) (Fig. 5D, O). The changes in *Ptgds* and *Hpgd* expression were far more significant and robust than the remaining differential gene expression in PD:ibot mice (Fig. 5A, B, D). L-PGDS synthesizes PGD2 extracellularly (Urade and Hayaishi, 2000) whereas Hpgd inactivates PGD2 (Conner et al., 2001). We hypothesize that an increase in L-PGDS combined with a decrease in Hpgd could lead to augmented extracellular PGD2. Blocking L-PGDS secretion and thus extracellular PGD2 synthesis could activate compensatory mechanisms to boost PGD2 by increasing the mRNA of L-PGDS (*Ptgds*) and decreasing the mRNA of *Hpgd*, which is consistent with criterion 3. Therefore, we next evaluated L-PGDS as a candidate molecule for the regulation of oligodendrocyte development.

### Blocking L-PGDS leads to oligodendrocyte development defects *in vitro*

We performed Western blot analyses to assess L-PGDS secretion by botulinum toxin B-expressing OPCs/oligodendrocytes in culture. We detected an increase in intracellular L-PGDS in OPC/oligodendrocyte cultures from PD:ibot mice compared with controls (Fig. 6A), consistent with the increase in *Ptgds* mRNA determined by RNA-seq (Fig. 5A, B, M, N). Secreted L-PGDS, however, is lower in PD:ibot compared with control cultures (Fig. 6A-C), suggesting that botulinum toxin B inhibits L-PGDS secretion. L-PGDS secretion is not completely eliminated, most likely because not all cells in the culture express botulinum toxin (efficiency: 60-80%, Fig. 1G) and wild-type cells may compensate by increasing secretion when extracellular L-PGDS levels are low.

**Figure 6.**
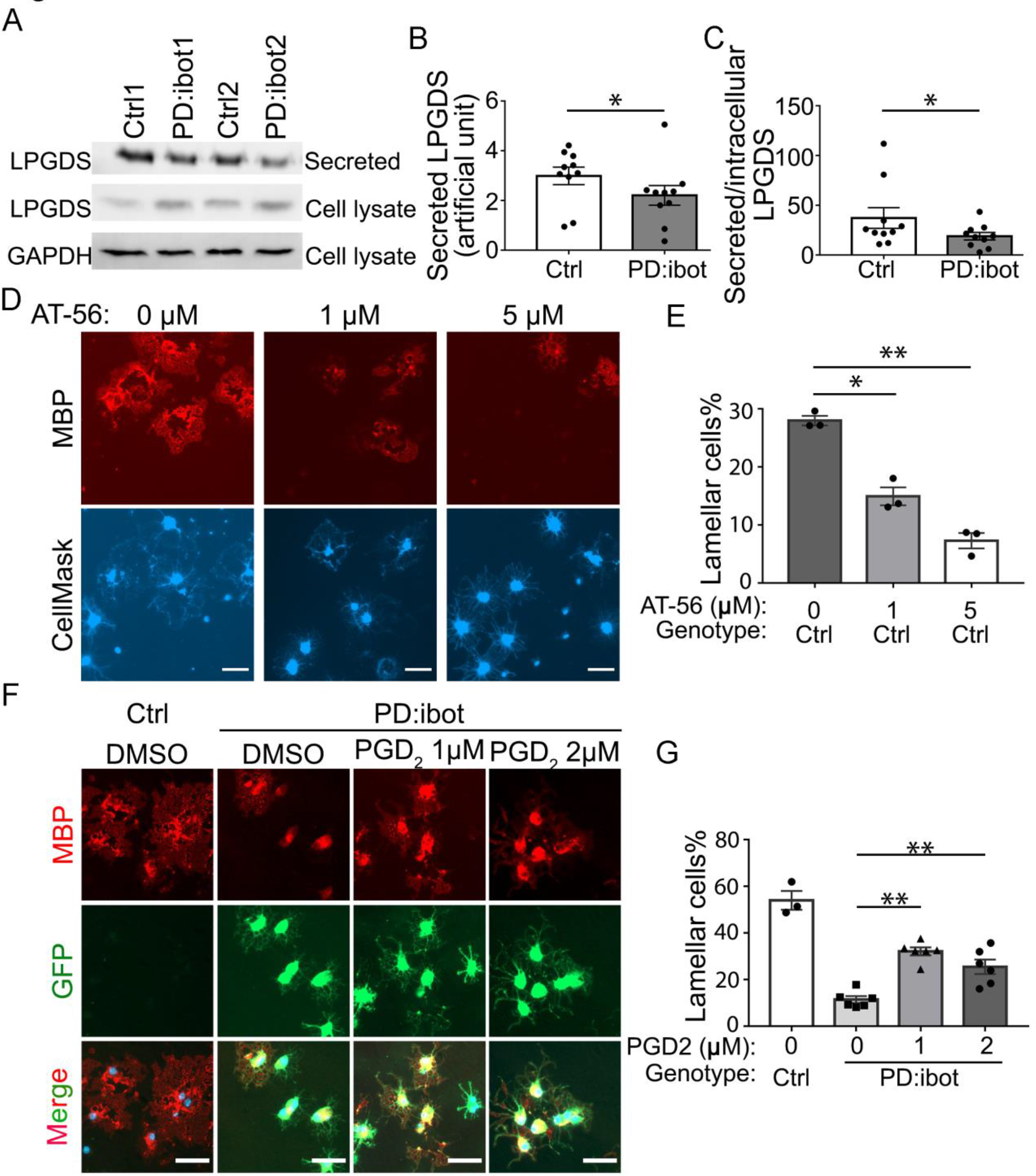
The effect of LPGDS inhibitor, AT-56, and PGD2 on oligodendrocyte development *in vitro*. (A) Immunoblot of secreted and intracellular L-PGDS protein from oligodendrocyte cultures from PD:ibot and littermate control mice. (B) Quantification of the immunoblot signal intensity of secreted L-PGDS. N=10 mice per group. Paired two-tailed T-test. (C) Quantification of the ratio of secreted and intracellular L-PGDS. N=10 mice per group. Paired two-tailed T-test. (D) Oligodendrocyte cultures from wild-type mice after 7 days of differentiation in the presence and absence of the LPGDS inhibitor AT-56. Red: MBP immunofluorescence. Blue: CellMask, which labels all cells. Scale bars: 40 μm. (E) Quantification of cells with lamellar morphology. One-way ANOVA with Dunnett’s test for multiple comparison correction. N=3 cultures from 3 mice per group. Lamellar cells%: DMSO control: 28±0.8; 1 μM AT-56: 15±1.6, p=0.036; 5 μM AT-56: 7.3±1.3, p=0.0050; (F) Partial rescue of oligodendrocyte differentiation by PGD2. Oligodendrocytes from PD:ibot and control mice in culture after 7 days of differentiation. Red: MBP immunofluorescence. Green: ibot-GFP, only present in cells from PD:ibot mice. Blue: DAPI labels nuclei of all cells. Scale bars: 50 μm. (G) Quantification of the percentage of lamellar cells. One-way ANOVA with Dunnett’s test for multiple comparison correction. N=3-6 cultures from 3 mice per group. PD:ibot + DMSO; 11.4±1.5; PD:ibot + 1 μM PGD2: 32.1±1.7, p=0.0001 compared with PD:ibot + DMSO; PD:ibot + 2 μM PGD2: 25.5±3.1, p=0.0013 compared with PD:ibot + DMSO; wild type control: 54±4, p=0.0001 compared with PD:ibot + DMSO.

To determine the role of L-PGDS in oligodendrocyte development and CNS myelination, we first assessed oligodendrocyte development *in vitro* in the presence of AT-56, a specific L-PGDS inhibitor (Irikura et al., 2009). We found that AT-56 inhibits wild-type oligodendrocyte development in a dose-dependent manner *in vitro* (Fig. 6D, E), suggesting a requirement of L-PGDS in oligodendrocyte development, without affecting their survival (Figure 6-figure supplement 1).

### L-PGDS is required for oligodendrocyte development and myelination *in vivo*

Having discovered the role of L-PGDS in oligodendrocyte development *in vitro*, we next assessed whether L-PGDS regulates oligodendrocyte development and myelination *in vivo*. We examined oligodendrocytes in L-PGDS global knockout mice and found a significant decrease in CC1^+^ oligodendrocytes and MBP^+^ myelin in the corpus callosum and cerebral cortex of L-PGDS-knockout mice at P9 (Fig. 7A-F). The density of OPCs (PDGFRα^+^) in the corpus callosum and cerebral cortex did not differ between L-PGDS- knockout and control mice (Fig. 7G, H).

**Figure 7.**
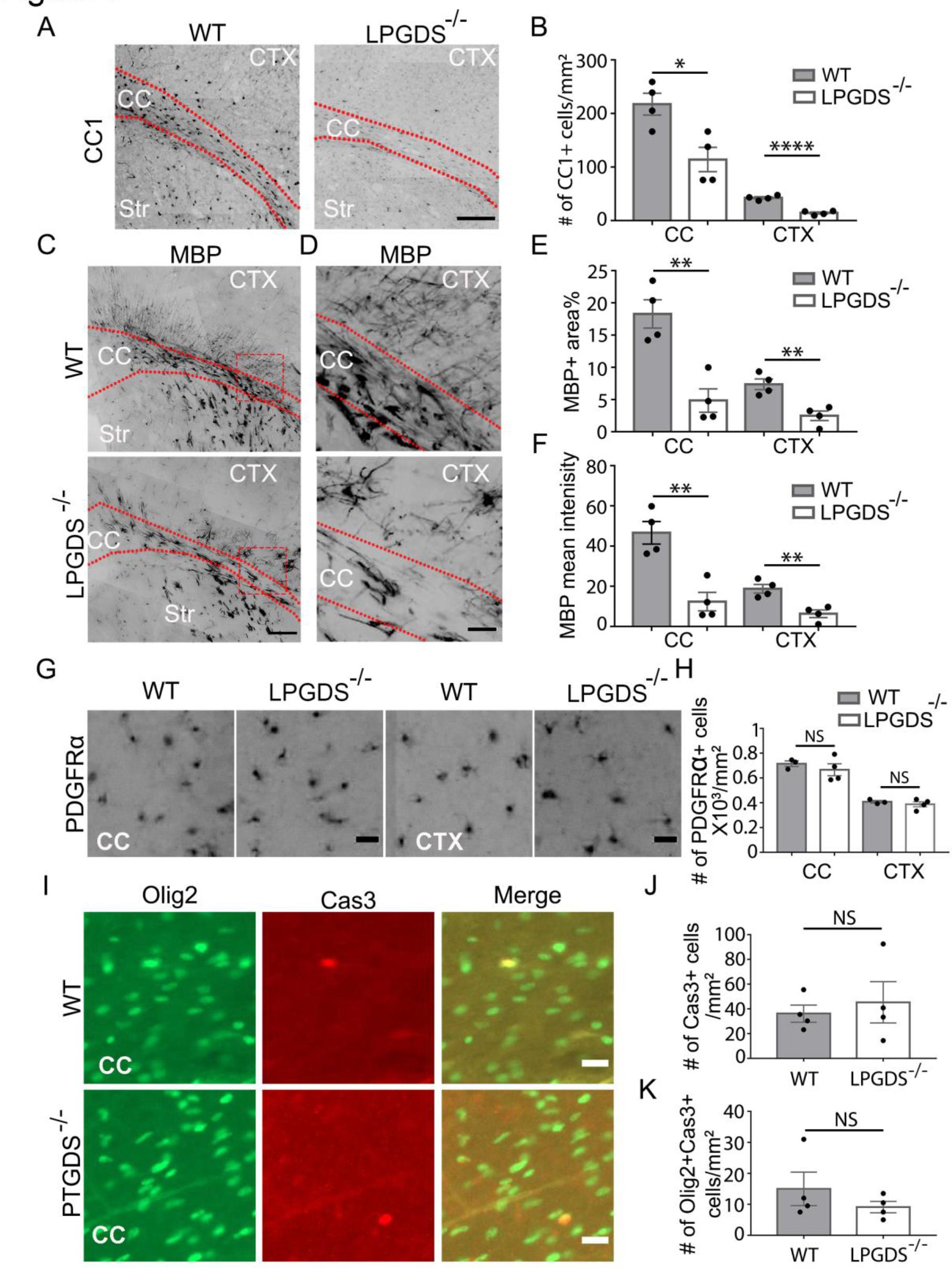
Oligodendrocyte and myelin protein defect in LPGDS-knockout mice. (A) CC1 immunofluorescence at P9. The dashed lines delineate the corpus callosum (CC). Ctx, cortex. Str, striatum. Scale bar: 200 μm. (B) Quantification of the density of CC1^+^ cells. N=4 mice per genotype. Corpus callosum: 217.6±20.3/mm^2^ in control, 114±22.8 in knockout, p=0.015; cerebral cortex: 42.5±2.2 in control, 14.3±2.0 in knockout, p=0.0001. Unpaired two-tailed T-test in all quantifications in this figure. (C) Myelin marker MBP immunofluorescence in the brains of LPGDS-knockout and littermate control mice at P9. Scale bar: 200 μm. (D) Enlarged view of the boxed areas in C. Scale bar: 50 μm. (E) Quantification of the percentage of MBP^+^ area. N=4 mice per genotype. Corpus callosum: 18.3±2.2 in control, 4.8±1.8 in knockout, p=0.0033; cortex: 7.4±0.8 in control, 2.5±0.7 in knockout, p=0.0046. (F) Quantification of the average MBP fluorescence intensities. Corpus callosum: 46.6±5.6 in control, 12.4±4.6 in knockout, p=0.0033; cortex: 18.8±2.1 in control, 6.3±1.9 in knockout, p=0.0046. (G) PDGFRα immunofluorescence at P9 in the corpus callosum and the cerebral cortex. Scale bar: 20 μm. (H) Quantification of the density of PDGFRα^+^ cells. Corpus callosum: 715.3±22.7 in control, 667±48.0 in knockout, p=0.46; cerebral cortex: 406.2±10.3 in control, 388.3±20.8 in knockout, p=0.48. (I) Activated caspase-3 immunofluorescence in LPGDS-knockout and littermate control mice. Scale bar: 20 μm. (J) Quantification of activated caspase-3^+^ cells. (K) Quantification of activated caspase-3^+^ oligodendrocyte-lineage cells.

### PGD2 restores the development of secretion-deficient oligodendrocytes

Next, we determined whether the product of the L-PGDS enzyme, PGD2, is sufficient to rescue the morphological maturation defect of PD:ibot cells *in vitro*. Indeed, exogenous addition of PGD2 partially restored the percentage of cells with lamellar morphology from PD:ibot mice (Fig. 6F, G), further supporting a role of L-PGDS in oligodendrocyte development. Although other cell types could mediate the effects of systemic L-PGDS knockout, the observation that PGD2 partially rescues the morphological maturation of oligodendrocytes in purified cultures *in vitro* supports a direct role of PGD2 in oligodendrocytes.

Given the enriched expression of L-PGDS by oligodendrocyte-lineage cells (Zhang et al., 2014, 2016) and its established localization in the extracellular space (Hoffmann et al., 1993), our results indicate that L-PGDS is an oligodendrocyte-lineage cell-secreted molecule that promotes oligodendrocyte development and myelination. Our results are consistent with the following model: OPCs secrete autocrine/paracrine signals such as L-PGDS to promote oligodendrocyte development and myelination. When VAMP1/2/3- dependent exocytosis is blocked, L-PGDS secretion is defective, and the cells upregulate the mRNA encoding L-PGDS (*Ptgds*) to compensate for the defect. Still, L-PGDS proteins cannot be released extracellularly, leading to defective oligodendrocyte development and myelination. L-PGDS-knockout mice exhibit similar defects. OPCs and oligodendrocytes secrete a variety of molecules. It is likely that other unidentified molecules also contribute to the oligodendrocyte-lineage cell-autocrine/paracrine loop. Nevertheless, our discovery of the role of L-PGDS in oligodendrocytes provides insight into the mechanisms regulating oligodendrocyte development and myelination.

## Discussion

In this study, we showed that oligodendrocyte-lineage cell-secreted molecules promote oligodendrocyte development and myelination in an autocrine/paracrine manner. We identified L-PGDS as one such secreted molecule, thus revealing a novel cellular mechanism regulating oligodendrocyte development.

Previously, the roles of VAMP3 and related pathways in myelin protein delivery and oligodendrocyte morphogenesis have been investigated largely *in vitro* using cultures of an OPC-like cell line (Oli-Neu cells) or primary oligodendrocytes. For example, VAMP3 and VAMP7 knockdown inhibits the transport of a myelin protein, Proteolipid Protein1 *in vitro* (Feldmann et al., 2011). Tetanus toxin, which cleaves VAMP1/2/3, inhibits oligodendrocyte branching *in vitro* (Sloane and Vartanian, 2007). Syntaxin4, a potential binding partner of VAMP3, is required for transcription of MBP in oligodendrocytes *in vitro* (Bijlard et al., 2015). Our study established a requirement of VAMP1/2/3-dependent exocytosis in oligodendrocyte development, myelination, and motor behavior *in vivo* and identified L-PGDS as an oligodendrocyte-lineage cell-secreted protein that promotes oligodendrocyte development and myelination.

VAMP1/2/3-dependent exocytosis is not the only pathway employed by oligodendrocyte-lineage cells to release molecules that mediate cell-cell interactions. For example, oligodendrocytes release exosome-like vesicles that inhibit the growth of myelin-like membranes *in vitro* (Bakhti et al., 2011). Tetanus toxin cleaves VAMP1/2/3 but does not affect exosome release (Fader et al., 2009). Therefore, the role of VAMP1/2/3-dependent exocytosis in promoting myelination and the effect of exosome-like vesicles in inhibiting myelination are likely parallel pathways independent of each other. In future studies, it could be interesting to determine the signals that regulate VAMP1/2/3-dependent exocytosis and VAMP1/2/3-independent exosome release during development and disease *in vivo* and thus define how these two seemingly opposing effects are coordinated to shape precise and dynamic myelination.

Our caspase-3 immunohistochemistry results did not show a difference in cell survival between PD:ibot and control cells, though we cannot exclude that cells die through non-apoptotic mechanisms or that microglia clear dying cells too rapidly for accurate counting.

OPCs are present throughout the CNS in adults (Hughes et al., 2013; Kang et al., 2010), even in demyelinated lesions in patients with multiple sclerosis (Franklin, 2002). Therefore, inducing oligodendrocyte and new myelin formation is an attractive strategy for treating demyelinating diseases. However, remyelination therapy has not been successful so far (Franklin, 2002), underscoring the need for a more complete understanding of the mechanisms regulating oligodendrocyte and myelin development. Our discovery of the role of L-PGDS in oligodendrocyte development and myelination adds to the knowledge of the molecular regulation of myelination. Interestingly, the *Ptgds* gene is lower in OPCs from multiple sclerosis patients than those from healthy controls (Jäkel et al., 2019). During remyelination in mice, PGD2 levels increase (Penkert et al., 2021). These observations and our results are consistent with the possible involvement of L-PGDS and PGD2 in remyelination. Future studies should investigate the role of L-PGDS in promoting remyelination.

A recent study shows that the gene encoding L-PGDS, *Ptgds*, marks a subpopulation of OPCs more resilient to spinal cord injury than other OPCs (Floriddia et al., 2020). Thus, the function of L-PGDS in OPC responses to injury and other neurological disorders will be interesting to explore in the future.

The product of the L-PGDS enzyme, PGD2, binds and activates two G-protein-coupled receptors, DP1 and DP2 (Gpr44) (Narumiya and Furuyashiki, 2011). In addition,

PGD2 undergoes non-enzymatic conversion to 15d-PGJ2, which activates the peroxisome proliferator-activated receptor-γ (Scher and Pillinger, 2005). Future studies should aim to identify the receptor(s) that mediates the effect of L-PGDS on oligodendrocyte development and myelination, as well as the downstream signaling pathways.

OPCs and oligodendrocytes secrete many molecules (Zhang et al., 2014). Although we identified the role of L-PGDS in oligodendrocyte development, our results do not rule out contributions from other secreted molecules. Our RNA-seq dataset provides a roadmap for future investigation of the roles of additional oligodendrocyte-lineage cell-secreted molecules in the brain.

Blocking exocytosis with botulinum toxin B may reduce the delivery of proteins and lipids to the plasma membrane, therefore causing cell-autonomous effects on oligodendrocyte development in addition to blocking secretion. Both cell-autonomous and cell-non-autonomous mechanisms may be involved in the effect of blocking exocytosis on oligodendrocyte development. Our insert rescue experiment (Fig. 4K-M) strongly supports the importance of secreted molecules but does not rule out cell-autonomous mechanisms. Investigating how the exocytosis pathway may contribute to oligodendrocyte development in a cell-autonomous manner may complement our study and improve the understanding of the role of VAMP1/2/3-dependent exocytosis in oligodendrocyte and myelin development in the future.

## Materials and Methods

### Lead contact and materials availability

Further information and requests for resources and reagents should be directed to and will be fulfilled by the Lead Contact, Ye Zhang. This study did not generate new unique reagents.

### Experimental animals

All animal experimental procedures were approved by the Chancellor’s Animal Research Committee at the University of California, Los Angeles, and conducted in compliance with national and state laws and policies. All the mice were group-housed in standard cages (maximum 5 mice per cage). Rooms were maintained on a 12-hour light/dark cycle. PDGFRα-CreER (Jax #018280) and ibot (Jax #018056) mice were obtained from Jackson Laboratories.

### OPC purification and culture

Whole brains excluding the olfactory bulbs and the cerebellum from one pup at postnatal day 7 to day 8 were used to make each batch of OPC culture. OPCs were purified using an immunopanning method described before (Emery and Dugas, 2013). Briefly, the brains were digested into single-cell suspensions using papain. Microglia and differentiated oligodendrocytes were depleted using anti-CD45 antibody- (BD Pharmingen, cat #550539) and GalC hybridoma-coated panning plates, respectively.

OPCs were then collected using an O4 hybridoma-coated panning plate. For most culture experiments, cells were plated on 24-well plates at a density of 30,000 per well. For comparison of OPC differentiation at different densities, OPCs were plated at densities of 5,000 per well and 40,000 per well. For all experiments, OPCs were first kept in proliferation medium containing growth factors PDGF (10 µg/ml, Peprotech, cat #100- 13A), CNTF (10 µg/ml, Peprotech, cat #450-13), and NT-3 (1 µg/ml, Peprotech, cat #450-03) for two to three days, and then switched to differentiation medium containing thyroid hormone (40 ng/ml, Sigma, cat #T6397-100MG) but without PDGF or NT-3 for seven days to differentiate them into oligodendrocytes as previously described (Emery and Dugas, 2013). Half of the culture media was replaced with fresh media every other day. All the cells were maintained in a humidified 37°C incubator with 10% CO2. Cells from both female and male mice were used. For coculture experiments with inserts, OPCs were purified from PD:ibot and littermate control mice as described above. 100,000 cells per well were plated on inserts with 1-μm diameter pores (VWR, cat #62406-173), and the inserts were placed on top of wells with cells plated at 40,000 cells per well density on 24-well culture plates. 200 μl medium was added per insert and 500 μl medium was added per well under the inserts.

### Drugs and treatment

4-hydroxy-tamoxifen stock solutions were made by dissolving 4-hydroxy- tamoxifen (Sigma, H7904) into pure ethanol at 10 mg/ml. The stock solutions were stored at -80°C until use. On the day of injection, an aliquot of 4-hydroxy-tamoxifen stock solution (100 μl) was thawed and mixed with 500 μl sunflower oil by vortexing for 5 min. Ethanol in the solution was vacuum evaporated in a desiccator (VWR, 89054-050) for an hour.

0.1 mg 4-hydroxy-tamoxifen was injected into each mouse subcutaneously daily for 2 days at P2 and P4. An L-PGDS inhibitor, AT-56 (Cayman Chemicals, cat #13160), and prostaglandin D2 (Cayman Chemicals, cat #12010) were dissolved in dimethyl sulfoxide (DMSO). To inhibit L-PGDS activity *in vitro*, AT-56 was added to the oligodendrocyte culture medium at 1 µM and 5 µM every other day. For prostaglandin D2 treatment, prostaglandin D2 was added to the oligodendrocyte culture medium at 1 µM and 2 µM every 12 hours. An equal amount of DMSO was added to the control wells. Because a metabolite of prostaglandin D2, 15-d-prostaglandin J2, induces cell death, which can be prevented by N-acetyl cysteine(Lee et al., 2008), we included 1 mM N-acetyl cysteine, which is shown to improve cell survival, in the culture media of prostaglandin D2-treated and control cells.

### RNA-seq

Total RNA was extracted using the miRNeasy Mini kit (Qiagen cat #217004). The concentrations and integrities of the RNA were measured using TapeStation (Agilent) and Qubit. cDNA was generated using the Nugen Ovation V2 kit (Nugen) and fragmented using the Covaris sonicator. Sequencing libraries were prepared using the Next Ultra RNA Library Prep kit (New England Biolabs) with 12 cycles of PCR amplification. An Illumina HiSeq 4000 sequencer was used to sequence these libraries and each sample had an average of 19.1 ± 2.9 million 50-bp single-end reads.

### RNA-seq data analysis

The STAR package was used to map reads to mouse genome mm10 and HTSEQ was used to obtain raw counts from sequencing reads. EdgeR-Limma-Voom packages in R were used to calculate Reads per Kilobase per Million Mapped Reads (RPKM) values from raw counts. The DESeq2 package was used to analyze differential gene expression.

### Immunohistochemistry and immunocytochemistry

Mice were anesthetized with isoflurane and transcardially perfused with phosphate-buffered saline (PBS) followed by 4% paraformaldehyde (PFA). The brains were removed and post-fixed in 4% PFA at 4°C overnight. The brains were washed with PBS and cryoprotected in 30% sucrose at 4°C for two days before embedding in optimal cutting temperature compound (Fisher, cat #23-730-571) and stored at -80°C. The brains were sectioned using a cryostat (Leica) into 30-μm-thick sections and floating sections were blocked and permeabilized in 5% donkey serum with 0.3% Tween-20 in PBS and then stained with primary antibodies against GFP (Aves Labs, Inc, cat #GFP-1020, dilution 1:500), PDGFRα (R&D Systems, cat #AF1062, dilution 1:500), Olig2 (Millipore, cat #211F1.1, dilution 1:500), CC1 (Millipore, cat #OP80, dilution 1:500), MBP (Abcam, cat #ab7349, dilution 1:500), and cleaved caspase-3 (Cell Signaling, cat #9661S, dilution 1:500) at 4°C overnight. Sections were washed three times with PBS and incubated with fluorescent secondary antibodies (Invitrogen) at 4°C overnight. Sections were mounted onto Superfrost Plus micro slides (Fisher, cat #12-550-15) and covered with mounting medium (Fisher, cat #H1400NB) and glass coverslips.

For immunocytochemistry of cultured cells, cells were fixed with 4% PFA and 0.3% Tween-20 in PBS. After blocking in 5% donkey serum, cells were then stained with the primary antibodies described above and BrdU antibodies (Abcam, cat #ab6326, 1:500) at 4°C overnight. After three washes in PBS, cells were stained with secondary antibodies and CellMask Blue (Invitrogen, cat #H32720, 1:1,000) at 4°C overnight. Cells were covered with mounting medium (Fisher, cat #H1400NB). The slides were imaged with a Zeiss Apotome epifluorescence microscope.

Fluorescence microscopy images were cropped and brightness contrast was adjusted with identical settings across genotype, treatment, and control groups using Photoshop and ImageJ. All the images were randomly renamed using the following website (https://www.random.org/) and quantified with the experimenter blinded to the genotype and treatment condition of the samples. Cells with MBP^+^ membrane spreading out were identified as lamellar cells. The illustrations were made with Biorender.

### Transmission electron microscopy

Brain specimens for transmission electron microscopy were prepared as described before (Salazar et al., 2018). Mice were anesthetized using isoflurane and transcardially perfused with 0.1M phosphate buffer (PB) followed by 4% PFA with 2.5% glutaraldehyde in 0.1M PB buffer. Brains were removed and post-fixed in 4% PFA with 2.5% glutaraldehyde in 0.1M PB for another two days. Brains were sliced with Young Mouse Brain Slicer Matrix (Zivic Instruments, cat #BSMYS001-1) and a small piece of the corpus callosum was isolated from brain sections at 0-1 mm anterior to Bregma. After wash, samples were then post-fixed in 1% osmium tetroxide in 0.1M PB (pH 7.4) and dehydrated through a graded series of ethanol concentrations. After infiltration with Eponate 12 resin, the samples were embedded in fresh Eponate 12 resin and polymerized at 60°C for 48 hours. Ultrathin sections of 70 nm thickness were prepared and placed on formvar/carbon-coated copper grids and stained with uranyl acetate and lead citrate. The grids were examined using a JEOL 100CX transmission electron microscope at 60 kV and images were captured by an AMT digital camera (Advanced Microscopy Techniques Corporation, model XR611) by the Electron Microscopy Core Facility, UCLA Brain Research Institute.

### Western blot

We purified OPCs from PD:ibot and control mice by immunopanning and cultured them in proliferation medium for 2-3 days and differentiation medium for 7 days as described above. To collect secreted samples, culture media were mixed with ethylenediaminetetraacetic acid (EDTA)-free protease inhibitor cocktail (Sigma, cat #4693159001) at a 6:1 ratio and centrifuged at 1000 × g for 10 min to remove dead cells and debris. To collect whole-cell lysates, cells were washed with PBS, lysed with radioimmunoprecipitation assay buffer containing EDTA-free protease inhibitor cocktail, and centrifuged at 12,000 × g for 10 min to remove cell debris.

All samples were mixed with sodium dodecyl sulfate (SDS) sample buffer (Fisher, cat # AAJ60660AC) and 2-mercaptoethanol before boiling for 5 min. Samples were separated by SDS-polyacrylamide gel electrophoresis, followed by transferring to polyvinylidene difluoride membranes via wet transfer at 300 mA for 1.5 hours. Membranes were blocked with clear milk blocking buffer (Fisher, cat #PI37587) for 1 hour at room temperature and incubated with primary antibodies against LPGDS (Santa Cruz Biotechnology, cat #sc-390717, dilution 1:1000), GAPDH (Sigma, cat #CB1001, dilution 1:5000), BoNT-B Light Chain (R&D Systems, cat #AF5420-SP, dilution 1:1000), VAMP2 (Synaptic Systems, cat #104 211, dilution 1:1000), and MBP (Abcam, cat #ab7349, dilution 1:1000) at 4°C overnight. Membranes were washed with tris-buffered saline with Tween 20 (TBST) three times and incubated with horseradish peroxidase-conjugated secondary antibodies (Mouse, Cell Signaling, cat #7076S; Rabbit, Cell Signaling, cat #7074S; Rat, Cell Signaling, cat #7077S; Sheep, Thermo Fisher, cat #A16041) for 1 hour at room temperature. After three washes in TBST buffer, SuperSignal™ West Femto Maximum Sensitivity Substrate (Fisher, cat #PI34095) was added to the membranes, and the signal was visualized using a ChemiDoc^TM^ MP Imaging system (BIO-RAD).

### Motor behavior

Mice were familiarized with being picked up and handled by the experimenter daily for three days before the test to reduce stress. Mice were also habituated to the rotarod testing room for 15 min prior to all testing. Both male and female adult mice (2 to 5 months old) were used in the rotarod test. Mice were given three trials per day for three consecutive days (5-60 rpm over 5 min, with approximately 30 min between successive trials). The latency to fall was measured and the experimenter was blinded to the genotype of the mice during the test.

### Quantification and statistical analysis

The numbers of animals and replicates are described in the figures and figure legends. The RNA-seq data were analyzed using the DESeq2 package. Adjusted P-values smaller than 0.05 were considered significant. For all non-RNA-seq data, analyses were conducted using Prism 8 software (Graphpad). The normality of data was tested by the Shapiro-Wilke test. For data with a normal distribution, Welch’s t-test was used for two-group comparisons and one-way ANOVA was used for multi-group comparisons. An estimate of variation in each group is indicated by the standard error of the mean (S.E.M.). * p<0.05, ** p<0.01, *** p<0.001. An appropriate sample size was determined when the study was being designed based on published studies with similar approaches and focus as our study. A biological replicate is defined as one mouse. Different culture wells from the same mouse or different images taken from the brains or cell cultures from the same mouse are defined as technical replicates. All statistical tests were performed with each biological replicate/mouse as an independent observation. The number of times each experiment was performed is indicated in figure legends. No data were excluded from the analyses. Mice and cell cultures were randomly assigned to treatment and control groups. Imaging analyses and behavior tests were conducted when the experimenter was blinded to the genotypes or treatment conditions.

### Data availability

We deposited all RNA-seq data to the Gene Expression Omnibus under accession number GSE168569. All relevant data are available from the authors without restrictions.

### Code availability

This study did not generate new codes.

## Acknowledgments

We thank Akiko Nishiyama, J. Bradley Zuchero, Hui Zong, and Mable Lam for their advice and editing of our manuscript. We thank Qingyun Li, Richard Breyer, Henry Lin, and Ginger Milne for their advice. We thank Garret FitzGerald for reagents. We thank the Eli and Edythe Broad Center of Regenerative Medicine and Stem Cell Research, UCLA BioSequencing Core Facility for their services, Mahnaz Akhavan and Suhua Feng for their technical support. This work is supported by the Knaub Postdoctoral Fellowship to L.P., the National Institute of Neurological Disorders and Stroke of the National Institute of Health (NIH) R00NS089780 and R01NS109025, the National Institute of Aging of the NIH R03AG065772, the National Center for Advancing Translational Science UCLA CTSI Grant UL1TR001881, the W. M. Keck Foundation Junior Faculty Award, the UCLA Eli and Edythe Broad Center of Regenerative Medicine and Stem Cell Research Innovation Award, the Ablon Faculty Scholar Award, and the Friends of the Semel Institute for Neuroscience & Human Behavior Friends Scholar Award to Y.Z.

## Author Contributions

(A) P. and Y.Z. conceived of the project and designed the experiments. L.P. performed most of the experiments. A.T. and L.P. analyzed the L-PGDS-knockout mice under the supervision of C.T. and Y.Z. L.O.S. contributed to experimental design. A.J.Z. assisted in mouse colony management and immunohistochemistry experiments. K.F., and Y.U. provided the L-PGDS-knockout mice. L.P. and Y.Z. analyzed the data and wrote the paper. All authors read the manuscript.

## Supplementary Information

### Supplementary Figures

**Figure 1-figure supplement 1.**
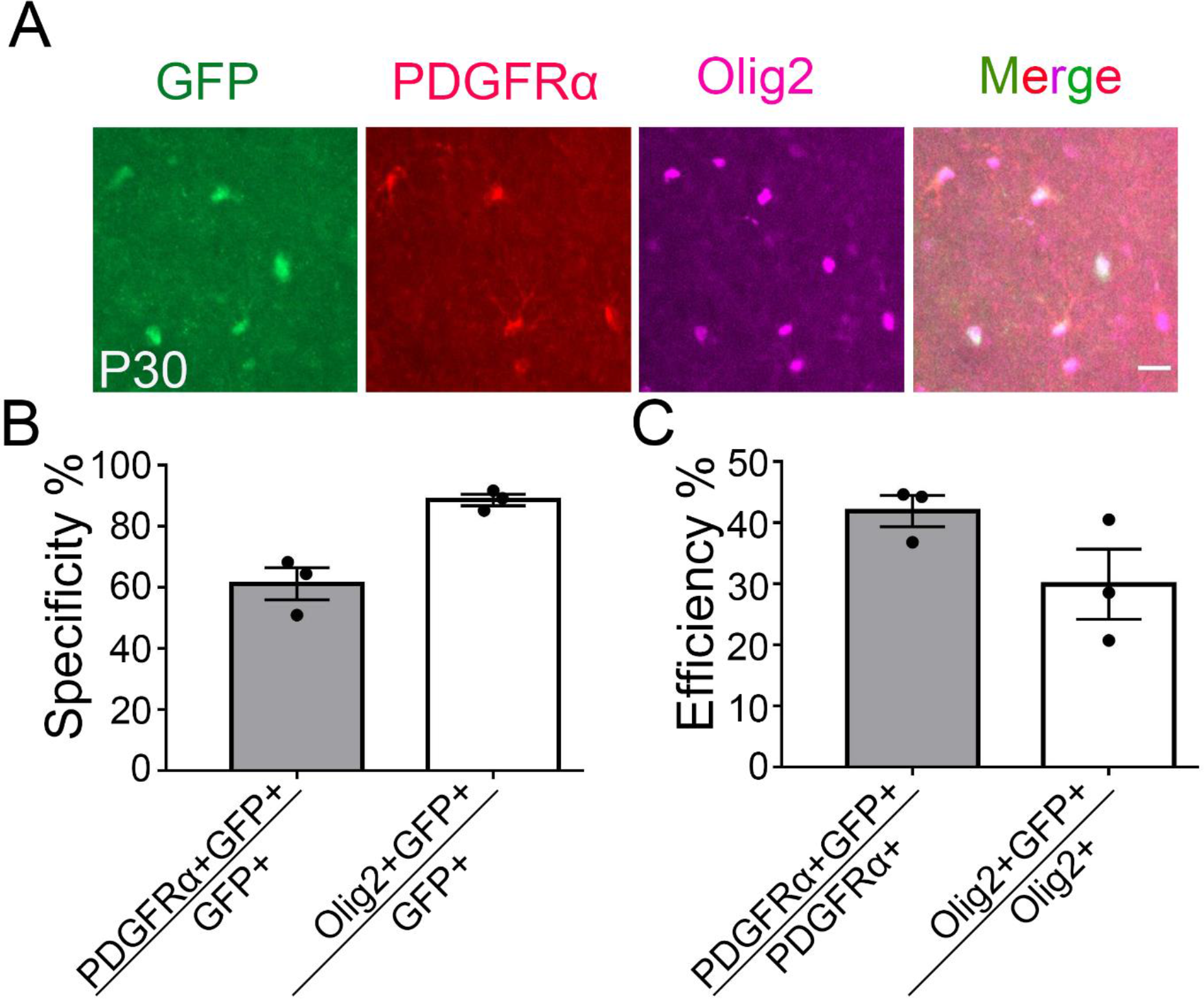
Specificity and efficiency of ibot expression in oligodendrocyte-lineage cells at P30. (A) Colocalization of ibot-GFP with PDGFRα and Olig2 in PDGFRα-CreER; ibot (PD:ibot) mice. Scale bar: 20 μm. (B) Specificity of ibot-GFP expression in oligodendrocyte-lineage cells at P30. N=3 mice per group. (C) Efficiency of ibot-GFP expression in oligodendrocyte-lineage cells at P30. N=3 mice per group.

**Figure 6-figure supplement 1.**
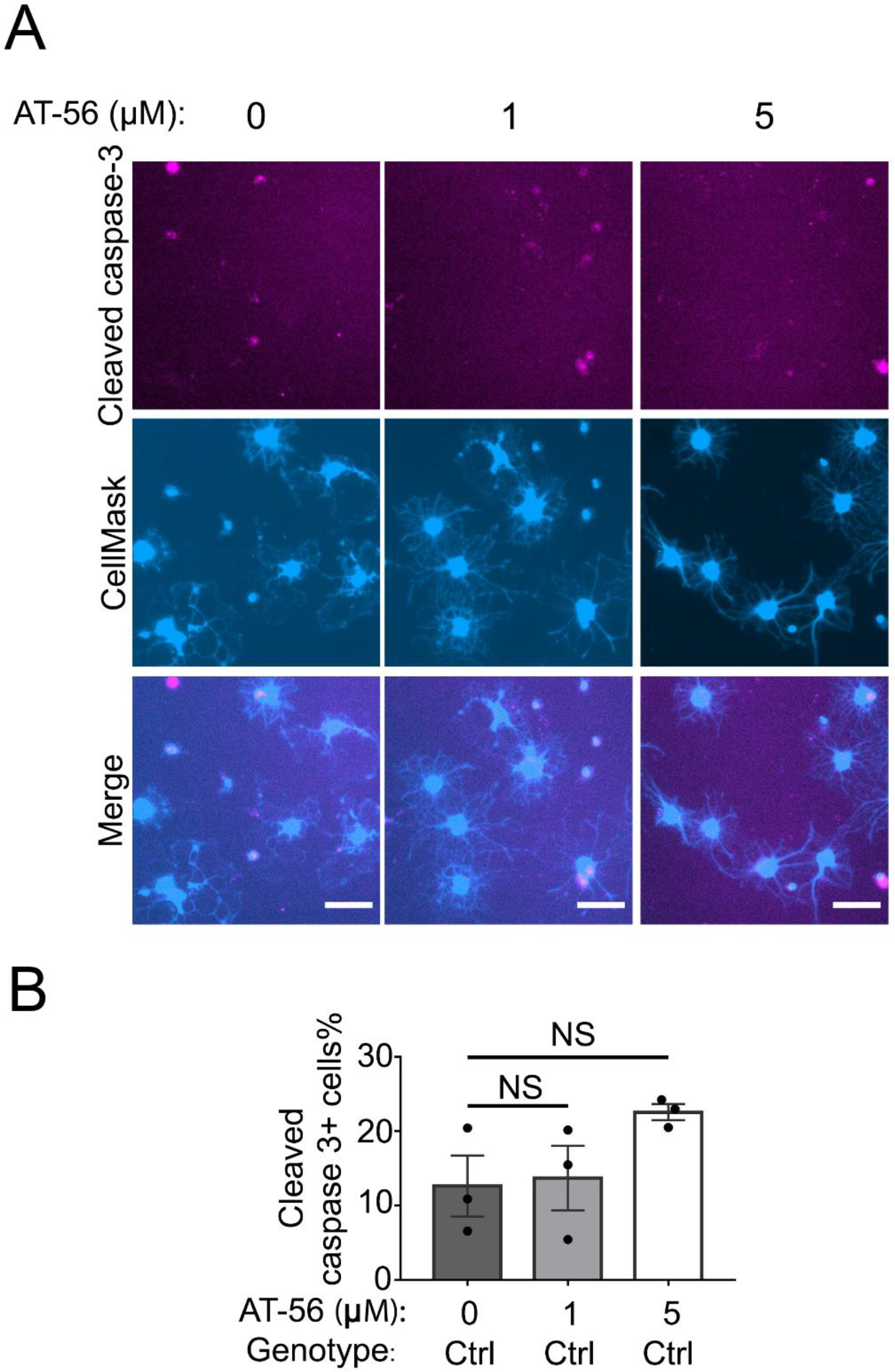
No change in the percentage of apoptotic cells in AT-56 treatment. (A) Oligodendrocyte culture from wildtype mice after 7 days of differentiation in the presence and absence of the LPGDS inhibitor AT-56. Magenta: cleaved caspase-3. Blue: CellMask, which labels all cells. Scale bars: 40 μm. (B) Quantification of caspase3^+^ cells. One-way ANOVA with Dunnett’s test for multiple comparison correction. N=3 cultures from 3 mice per group.

### Supplementary Tables

Supplementary Table 1. Gene expression (RPKM) of OPCs, oligodendrocytes, microglia, and astrocytes from PD:ibot and littermate control mice at P17 determined by RNA-seq Reads per kilobase of transcripts per million mapped reads (RPKM) are shown.

Supplementary Table 2. Differentially expressed genes in OPCs, oligodendrocytes, microglia, and astrocytes from PD:ibot and littermate control mice at P17 Genes with adjusted P-values <0.05 are shown. We used DESeq2 to determine differential gene expression.

Supplementary Table 3. Gene ontology terms associated with differentially expressed genes in OPCs, oligodendrocytes, microglia, and astrocytes from PD:ibot and littermate control mice at P17 No gene ontology terms were significantly enriched in genes downregulated in OPCs, astrocytes, or upregulated in microglia in PD:ibot mice.

